# SAINT: automatic taxonomy embedding and categorization by Siamese triplet network

**DOI:** 10.1101/2021.01.20.426920

**Authors:** Yang Young Lu, Yiwen Wang, Fang Zhang, Jiaxing Bai, Ying Wang

## Abstract

**Motivation:** Understanding the phylogenetic relationship among organisms is the key in contemporary evolutionary study and sequence analysis is the workhorse towards this goal. Conventional approaches to sequence analysis are based on sequence alignment, which is neither scalable to large-scale datasets due to computational inefficiency nor adaptive to next-generation sequencing (NGS) data. Alignment-free approaches are typically used as computationally effective alternatives yet still suffering the high demand of memory consumption. One desirable sequence comparison method at large-scale requires succinctly-organized sequence data management, as well as prompt sequence retrieval given a never-before-seen sequence as query.

**Results:** In this paper, we proposed a novel approach, referred to as SAINT, for efficient and accurate alignment-free sequence comparison. Compared to existing alignment-free sequence comparison methods, SAINT offers advantages in two aspects: (1) SAINT is a weakly-supervised learning method where the embedding function is learned automatically from the easily-acquired data; (2) SAINT utilizes the non-linear deep learning-based model which potentially better captures the complicated relationship among genome sequences. We have applied SAINT to real-world datasets to demonstrate its empirical utility, both qualitatively and quantitatively. Considering the extensive applicability of alignment-free sequence comparison methods, we expect SAINT to motivate a more extensive set of applications in sequence comparison at large scale.

**Availability:** The open source, Apache licensed, python-implemented code will be available upon acceptance.

**Supplementary information:** Supplementary data are available at *Bioinformatics* online.

## 1 Introduction

Understanding the phylogenetic relationship among organisms is the key in contemporary evolutionary study (Delsuc *et al*., 2005). Owing to the rapid developments in sequencing technologies, biologists nowadays have been able to collect complete or near-complete genome sequences, without relying on cultivation or reference genomes (Hug *et al*., 2016). These readily available genome sequences are phylogenetically informative to classify organisms and as a consequence, make the sequence-based comparison methods popular and widely applicable to study the phylogenetic relationship among a set of organisms (Eisen and Fraser, 2003; Bernard *et al*., 2019).

Due to the availability of complete or near-complete genome sequences, it is straightforward to identify homologous regions that are shared among different organisms by using in a multiple sequence alignment (MSA) (Feng and Doolittle, 1987). The workhorse for MSA, sequence alignment (Altschul *et al*., 1990), can be either global or local, where global alignment focuses on the entirety (Needleman and Wunsch, 1970) whereas local alignment only highlights local regions of high similarity (Smith *et al*., 1981). Despite wide applicability and great popularity, alignment-based sequence comparison methods are not appropriate in some situations. First, alignment based methods are not only computationally expensive, not also memory intensive in constructing large-size indexing files, thus not scalable to large-scale sequence data. Second, genomes of bacteria often contain DNA stretches acquired either from unrelated organisms or from lateral gene transfer (Bernard *et al*., 2019). However, alignment based methods are usually incapable of distinguishing those lateral signals. And finally, with the advent of next-generation sequencing (NGS) technologies, unprecedented high volumes of short reads have been generated which is very challenging to assemble. Therefore, it is desirable yet infeasible for alignment-based methods to compare genomes from unassembled sequence reads.

Alignment-free sequence comparison methods (Lu *et al*., 2017; Zielezinski *et al*., 2017, 2019; Lu *et al*., 2020), as attractive alternatives, are more resilient to complicated genetic rearrangements and much more computationally efficient than conventional alignment-based methods for studying the relationships among sequences. In brief, most alignment-free methods compare the sequences in terms of the statistics of fixed-length word (also known as *k*-mer) frequency. Specifically, the vectors comprised of the composition of *k*-mers, usually referred to as *k*-mer frequency vectors, need to be computed for comparison. The underlying rationale of using *k*-mer is based upon the assumption that *k*-mer patterns shared across a set of sequences can potentially capture the phylogeny informative homology signals. However, the *k*-mer frequency vectors for large k of practical interest demand excessive memory and storage consumption, undermining the popularity of alignment-free methods in large scale. Furthermore, existing alignment-free methods are all data-independent in the sense that dissimilarity measures are predefined in a heuristic manner and thus only provide rough approximation of the proximity among organisms (Zheng *et al*., 2019).

Here, considering the comparison within a massive collection of genomic sequences, one desirable sequence comparison method at this scale requires succinctly-organized sequence data management, as well as prompt sequence retrieval given a never-before-seen sequence as query. In this paper, we propose a new method, referred to as SAINT (<u>SiA</u>mese trIplet <u>N</u>etwork for <u>T</u>axonomy embedding and categorization), for efficient and accurate alignment-free sequence comparison (Figure 1). The key idea of SAINT is: (1) break down a phylogenetic tree containing the interspecies relationships among different organisms by the relative triplet comparison in the form of “A is closer to B than C”; (2) use a specific deep neural network to learn an embedding function for sequence triplets so that in the embedding space, a small distance between similar sequence pair A and B is favoured whereas dissimilar sequence pair A and C is pushed apart by a large distance.

**Fig 1.**
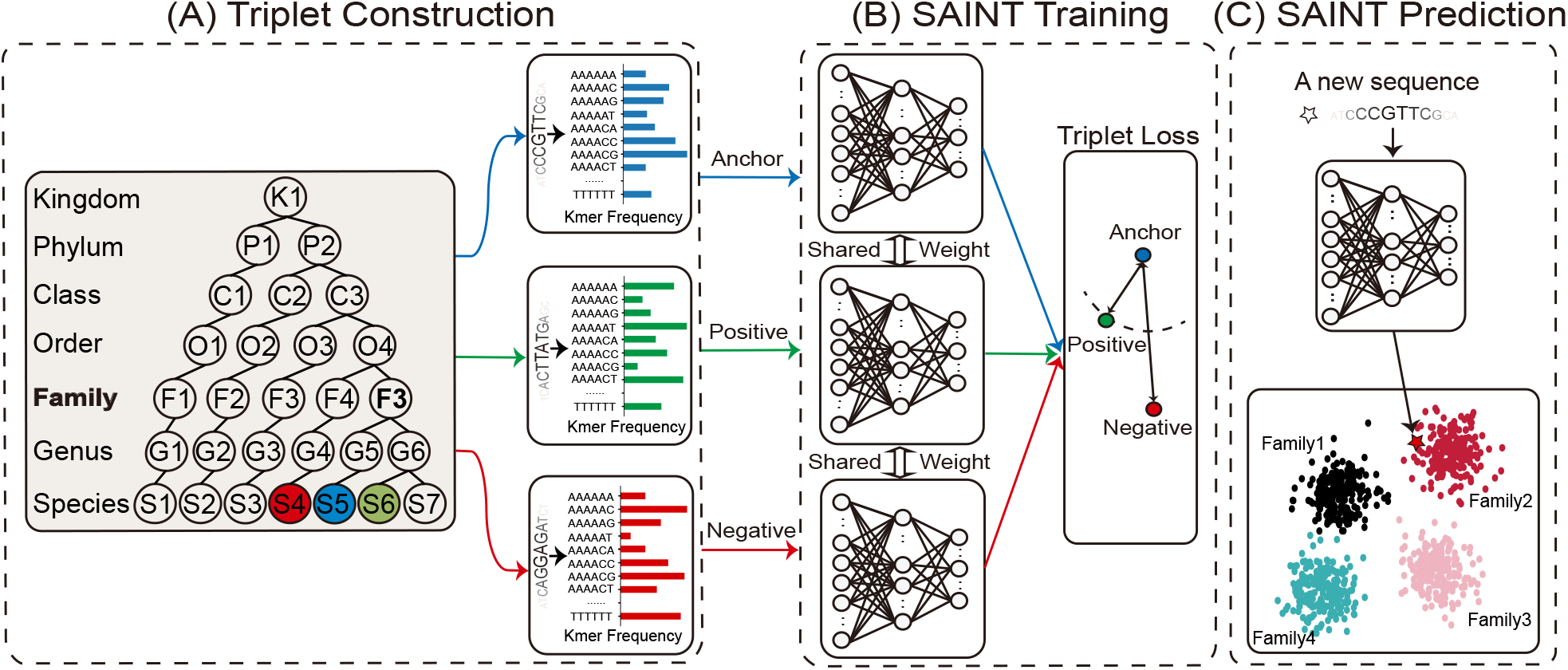
The workflow of SAINT. (A) SAINT encodes the phylogeny into a set of sequence triplets, each of which is represented as a *k*-mer frequency vector; (B) Each triplet, comprised of anchor, positive and negative sequences, is passed through a Siamese triplet network as input, and ultimately embedded into a common latent space for each sequence. The contrastive loss function penalizes far-apart anchor and positive sequences as well as nearby anchor and negative sequences, up to a pre-specified margin; (C) An unknown sequence is passed through the trained Siamese triplet network, projected into the embedding space among all trained sequences, and categorized by matching to the nearest neighbor.

Compared to existing alignment-free sequence comparison methods, SAINT offers following advantages:

- SAINT is more computationally fast and memory efficient because sequence data are operated in a compressed embedding space which is much faster to retrieval and succinct to store.
- SAINT is a weakly-supervised learning method where the embedding function is learned automatically from the easily-acquired data. Compared to existing deep learning-based alignment-free method (Zheng *et al*., 2019), SAINT doesn’t require tedious labors to collect accurate alignment distances to train.

We also apply SAINT to real-world datasets to demonstrate its empirical utility, both qualitatively and quantitatively. Considering the extensive applicability of alignment-free sequence comparison methods, we expect SAINT to motivate a more extensive set of applications in sequence comparison at large scale

## 2 Methods

### 2.1 Represent sequence by *k*-mer frequencies

The *k*-mer representation is prevalently used in alignment-free methods (Lu *et al*., 2017; Zielezinski *et al*., 2017), serving as the basis for genomic and metagenomic comparisons (Bernard *et al*., 2018; Wang *et al*., 2014). Concretely, given an alphabet Σ = {*A, C, G, T*} as well as a pre-specified number *k*, these alignment-free methods first construct a dictionary consisting of all possible subsequences of length *k*, then project the input sequence into a feature vector where each entry encodes the occurrence of the corresponding *k*-mer detected in the sequence. It is worth mentioning that there exists several variants of such feature vector, one commonly-used alternative, often referred to as the frequency vector, is to normalize the *k*-mer counts into frequencies by dividing the total counts of all *k*-mers; another popular alternative, often referred to as the indicator vector, is to binarize the *k*-mer counts into the presence/absence of *k*-mers. Since some previous studies suggested that using *k*-mer frequencies is more informative and generalizable than the presence/absence of k-mers (Lu *et al*., 2020), SAINT uses the *k*-mer frequencies as the sequence feature throughout this paper.

### 2.2 Encode phylogeny into triplets

SAINT encodes a phylogenetic tree containing the interspecies relationships among different organisms by the relative triplet comparison in the form of “A is closer to B than C”. For example, two genome sequences belonging to the same genus taxonomic level is more likely to be categorized into the same operational taxonomic unit (OTU) than those in higher genus levels such as family, order, class, etc. It is worth noting that such triplet encoding is equally informative as a phylogenetic tree in the sense that the phylogenetic tree can be accurately reconstructed based upon the triplets (Ranwez *et al*., 2010).

Mathematically, we represent a triplet by *t* = (*x, x*^+^, *x*^−^) where *x, x*^+^, and *x*^−^ are the feature vectors of the anchor sequence, positive sequence and negative sequence, respectively. The anchor and positive sequences belong to the same taxonomic level, whereas the negative sequence belongs to a different taxonomic level.

#### 2.2.1 Triplet sampling

Given a phylogeny with N sequences, the number of possible triplets is on the order of 𝒪 (*n*^3^). When constructing triplets from a large phylogeny, it is computationally unfeasible to train a model on such a prohibitively large number of triplets. Therefore, in practice it is often desirable to select a small representative subset of triplets through some sampling strategy.

One straightforward strategy is *k*-negative sampling strategy (Wu *et al*., 2017). In particular, each anchor and positive sequence pair (*x, x*^+^) is first selected, followed by *k* randomly selected negative sequences. However, it is not only challenging to determine a proper data-dependent *k* but also computational intensive to take excessive random selections into consideration. Another alternative sampling strategy is to select triplets that nearly violate the triplet constraints (Le Capitaine, 2018). In other words, given the anchor sequence *x*, the selected positive sequence *x*^+^should be furthermost from the anchor in the positive space whereas the negative sequence *x*^−^ should be closest to the anchor among all possible negatives. However, solely restricting to those difficult triplets may cause the training procedure unstable to converge.

In this paper, SAINT follows a combined sampling strategy shown as follows. Specifically, by firstly specifying the anchor sequence *x* and a target taxonomic level we select the positive sequence *x*^+^ such that the lowest common taxonomical ancestor (LCA) (Huson *et al*., 2007) between *x* and *x*^+^ matches the target taxonomic level. Afterwards, SAINT selects the negative sequence *x*^−^ with respective to different *classification difficulty* by enumerating the LCA between *x* and *x*^−^ from the immediate ancestor of the target taxonomic level to the most distal one. For example, as shown in Figure 1(A), by specifying the anchor and positive sequence S5 and S6 with the taxonomic level as Family, SAINT constructs four triplets by selecting four negative sequences S4, S3, S2, and S1 corresponding to the LCA as Order, Class, Phylum, and Kingdom, respectively.

### 2.3 Project triplets into a common embedding space

SAINT aims to learn a mapping function 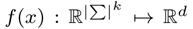, where |Σ|^*k*^ indicates the size of the *k*-mer frequency vector and *d* denotes the dimensionality of common embedding space. Given a triplet 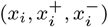 for *i* = 1, 2,…, *T* where *T* is the total number of the triplets, the learned function *f* (*x*) is expected to jointly pull similar sequence pair *x*_*i*_ and 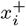 closer and push the dissimilar sequence pair *x*_*i*_ and 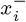 apart, in terms of the Euclidean distance. The key to the learning process is the contrastive loss function, defined as:

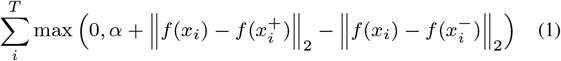

where ∥ · ∥_2_ indicates the *L*_1_-norm and *α >* 0 is a margin to separate the positive and negative pairs. Intuitively, Equation 1 penalizes far-apart anchor and positive sequences as well as nearby anchor and negative sequences, up to the pre-specified margin *α*. In this paper we fix *α* = 0.5.

The heart of learning the mapping function *f* (*x*) subject to the contrastive loss is the “Siamese triplet network” (Hoffer and Ailon, 2015), as illustrated in Figure 1(B). Specifically, the Siamese triplet network takes triplet sequences as input, and feed them independently into three identical embedding networks operated side by side. The embedding network parameters are shared by the three networks so that updates to one embedding network are always reflected in others. The shared parameters are learned by minimizing the contrastive loss function defined in Equation 1. Thus we allow the back-propagation from this loss function to update the weights in the network with regard to all triplet sequences simultaneously.

#### 2.3.1 Network implementation

In our implementation, the embedding network is represented by a fully connected multilayer perceptron (MLP), of which the number of nodes in the input layer equals the size of the *k*-mer frequency vector, *i*.*e*., |Σ|^*k*^. The MLP network has multiple alternating linear transformation and nonlinear activation layers. Each layer learns a mapping from its input to a hidden space, and finally, the last layer learns a mapping directly from the hidden space to the common embedding space of dimensionality *d*. In this work, we use an MLP with 6 hidden layers, each containing 200 neurons with ReLU activation. And the size of the output layer (*i*.*e*., the embedding dimensionality *d*) is set to be *d* = 100. We use Adam Kingma and Ba (2014) to train the model using an initial learning rate of 10^−4^ and batch size 5000. See Section 3.4 for the effect of different embedding dimensionality settings.

#### 2.3.2 Triplet Weighting

Equation 1 assumes that all triplets are important, which may not be ideal in model training. In particular, the contrastive loss, can be easily fulfilled in the cases where the negative sequence is significantly dissimilar to the anchor (*e*.*g*., the anchor and the negative sequence belong to different Class), but fails to discriminate the cases where the negative sequence is more or less similar to the anchor (*e*.*g*., the anchor and the negative sequence belong to the same Family but different Genus). In other words, attentions need to be paid on cases that are difficult to discriminate. In this paper, instead of assigning identical weights, we propose to assign a triplet-specific weight for each of the triplets so that difficult triplets are weighed more highly than easy ones. More precisely, we adapt the contrastive loss function in Equation 1 as follows:

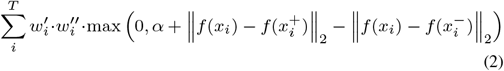

where 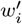 adjusts the *i*-th triplet inversely proportional to the frequencies of its corresponding classification difficulty level in the training data. And 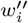 highlights the importance of the *i*-th triplet with respect to its classification difficulty level. Specifically, by letting *d*_*i*_ denote the LCA gap between 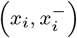 and 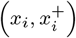, we define 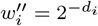. For example, as shown in Figure 1(A), the LCA gap between Family and Order is 1.

### 2.4 Categorize an unknown species by the phylogeny embeddings

SAINT is able to automatically categorize an unseen sequence into a trained phylogeny. To achieve this, briefly speaking, given an unseen sequence *x* and a targeted taxonomy level *t* (*e*.*g*., Genus, Family, Order, etc.), SAINT first infers the embedding distances between x and all sequences in the phylogeny, and then locates the best matching taxonomic unit by averaging the taxonomy-specific sequences. Specifically, by denoting the phylogeny sequence set as 𝒳 and its possible taxonomic units at the targeted taxonomy level as 𝒞_*t*_ = {*x*_*i*_ ∈ 𝒳: T(*x*_*i*_, *t*)} where T(*x*_*i*_, *t*) returns the corresponding taxonomic unit of *x*_*i*_ at the targeted taxonomy level *t*. SAINT aims to identify the best match in 𝒞_*t*_:

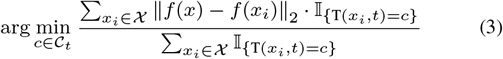

where 𝕀_{·}_ refers to an indicator function.

## 3 Results

### 3.1 Datasets

In the experiment, we constructed two genomic datasets downloaded from NCBI RefSeq Database (Pruitt *et al*., 2005). Both datasets, comprised of 1239 bacteria and 137 vertebrate sequences respectively, have detailed taxonomic hierarchy from Species, Genus, up to Phylum. We observe that, in each of the two datasets, when we go down the taxonomic hierarchy, the number of distinct taxonomic units increases notably (Figure 2). Accordingly, we adapt this observation in the training and testing division. Specifically, we split the training and testing sets in a way not only the taxonomic units under each taxonomic level are evenly split but each taxonomic unit appearing in the testing set is ensured to be included in the training set as well. We used the sampling method introduced in Section 2.2.1 to generate 11200 triplets in our training set and 10026 in our testing set for the vertebrate dataset, and 896240 triplets in both training and testing sets for the bacteria dataset, respectively. Within the training set, we further applied a 2-fold splitting scheme to evenly split the training and validation triplets.

**Fig 2.**
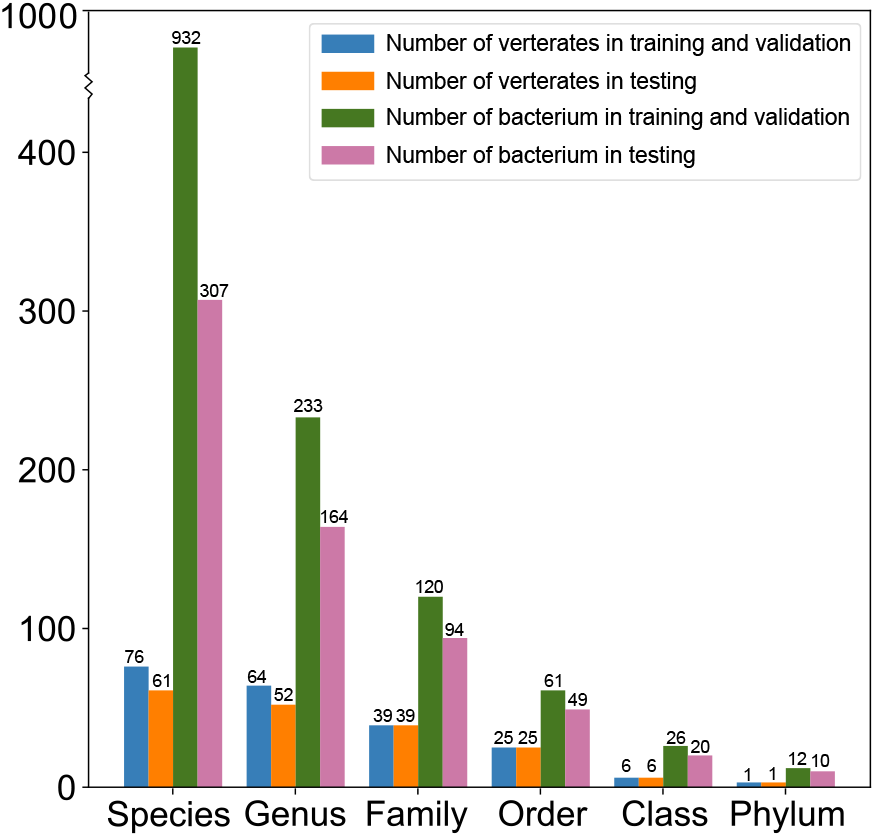
The summary of the training and testing sets splitting on vertebrate and bacteria datasets.

### 3.2 Performance on triplets classification

We first investigated whether the SAINT model is well-trained and properly behaved as expected. As shown in Figure 3(D) and 3(I), the contrastive loss function in Equation 1 for the training set drops sharply during the first dozens epochs before reaching a plateau, aligned with the trend for the testing set. In addition to the loss in general, we further scrutinized the actual learned relationship of triplets. As shown in both Figure 3(A)-(C) for the bacteria dataset and Figure 3(F)-(H) for the vertebrate dataset, the learned distances betweens the anchor and positive sequences are consistently smaller than the ones between the anchor and negative sequences. In short, SAINT successfully learned the embedding for all genome sequences which are not only capable of pulling the sequence pairs closer and pushing the dissimilar pairs apart, but also generalizable to sequences never seen before.

**Fig 3.**
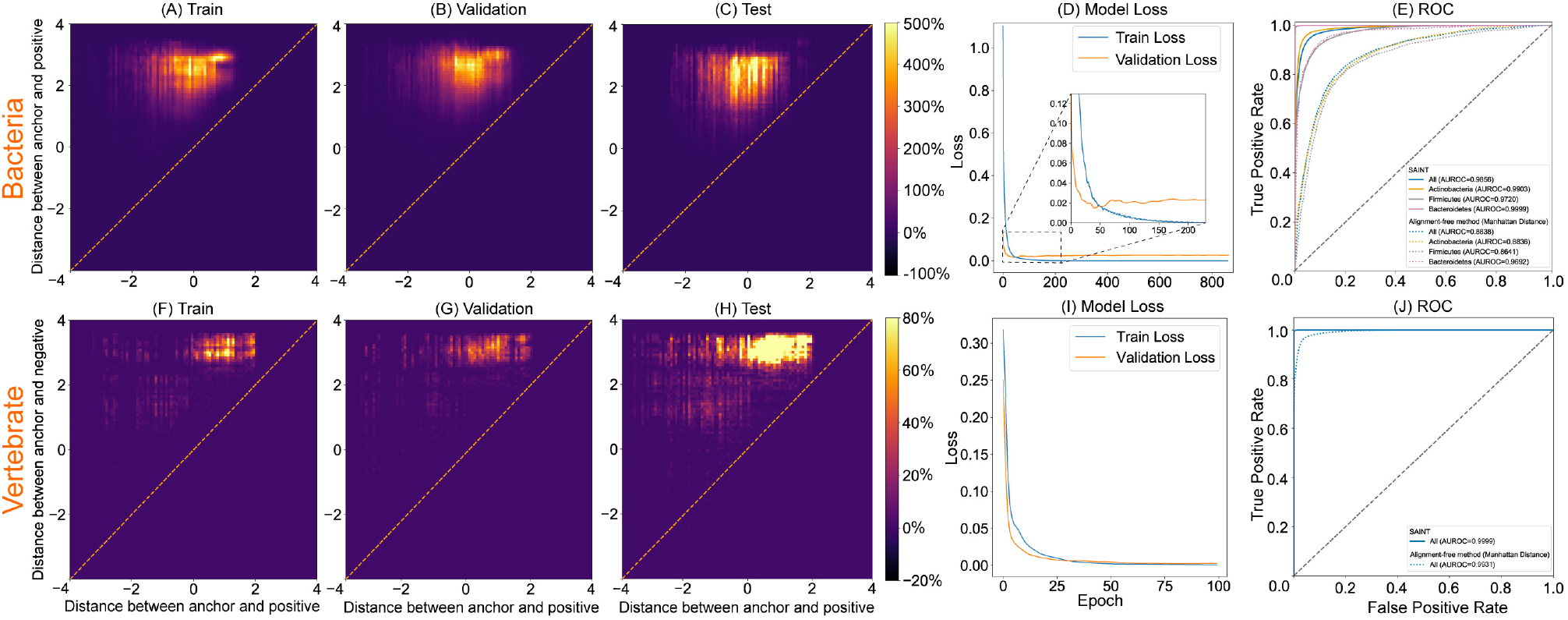
The performance of SAINT in the triplet classification task on bacteria and vertebrate datasets, depicted by (A)-(E) and (F)-(J), respectively.

We next evaluated SAINT in the context of triplet classification on both datasets. We notice that the triplets in the testing set are valid, that is, with the positive labels exclusively. Hence, we randomly chose half triplets in the testing set as the negative set by swapping the corresponding positive sequence and negative sequence. For each triplet 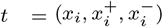, the prediction score is measured by 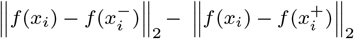, where the large value indicates that the distance between *x*_*i*_ and 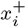 is much smaller than the one between *x*_*i*_ and 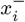. Measured by the area under the receiver operating characteristic curves (AUC), SAINT performs substantially better than a widely-used alignment-free dissimilarity metric, Manhattan distance, by an average increase of 10.18% on the bacteria dataset. For the vertebrate dataset, despite the fact that the performance of Manhattan distance is already as good as 0.99 in terms of AUC score, SAINT still improve the performance by reaching 1.00. Furthermore, for the bacteria dataset, we examined the performance of SAINT in three most common Phylums of bacteria, Actinobacteria, Firmicutes, and Bacteroidetes, SAINT remains the notable superiority in terms of AUC over Manhattan distance, by an average increase of 9.17%.

### 3.3 Performance on taxonomy localization

In this section, we systematically evaluated the prediction accuracy of SAINT in terms of each taxonomic level on both datasets. To alleviate the intrinsic stochasticity brought from neural network training, we thus repeated the 2-fold splitting on the training dataset for 6 times and reported the averaged performance. For the bacteria dataset shown in Figure 4(A), the taxonomic accuracy of SAINT reaches 0.84, 0.93, 0.95, 0.97, and 0.98 with respect to the level of Genus, Family, Order, Class, and Phylum, respectively. In comparison, the taxonomic accuracy of Manhattan distance only achieves 0.83, 0.87, 0.90, 0.94, and 0.96 correspondingly, with an average decrease of 3.88%. Analogously, for the vertebrate dataset shown in Figure 5(A), the taxonomic accuracy obtained by SAINT is 0.90, 0.97, 1.00, and 1.00 with respect to the level of Family, Order, Class, and Phylum, respectively, achieving an average increase of 4.60% compared to Manhattan distance. It is worth mentioning that the Genus level comparison is not shown because there are too few samples of each category in the genus layer.

**Fig 4.**
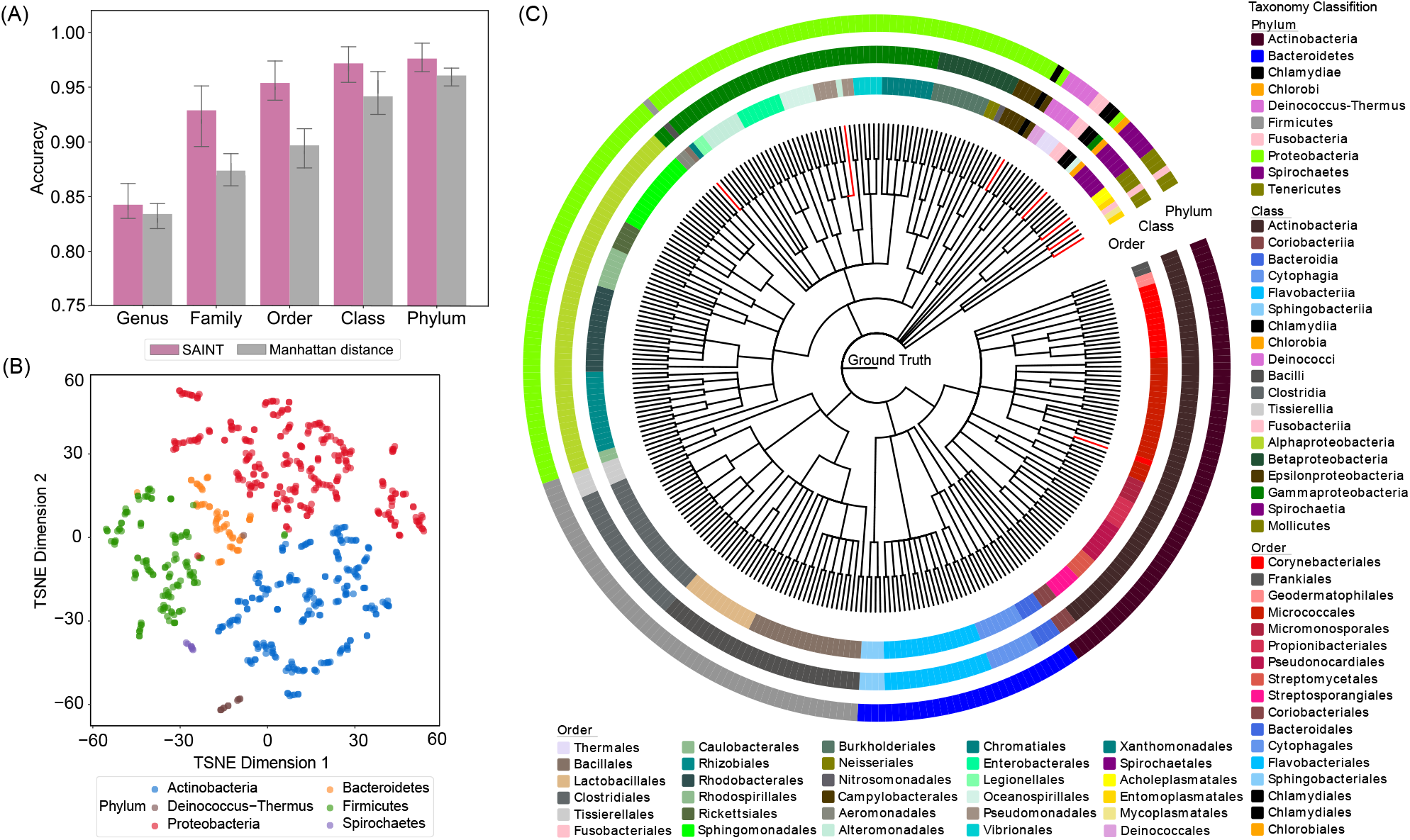
The performance of SAINT in the taxonomy localization task on bacteria dataset. (A) The prediction accuracy of SAINT and Manhattan distance in terms of different taxonomic level from Genus up to Phylum; (B) The embedding representation learned by SAINT can separate different common bacterial phylums into distinct groups by using t-SNE visualization; (C) The predicted triplets which encode 307 bacteria reference genomes in the testing set are accurately aligned to the true phytogenetic tree downloaded from NCBI. Note that the branches marked in red indicate the inconsistency between the prediction and the reference phytogeny.

**Fig 5.**
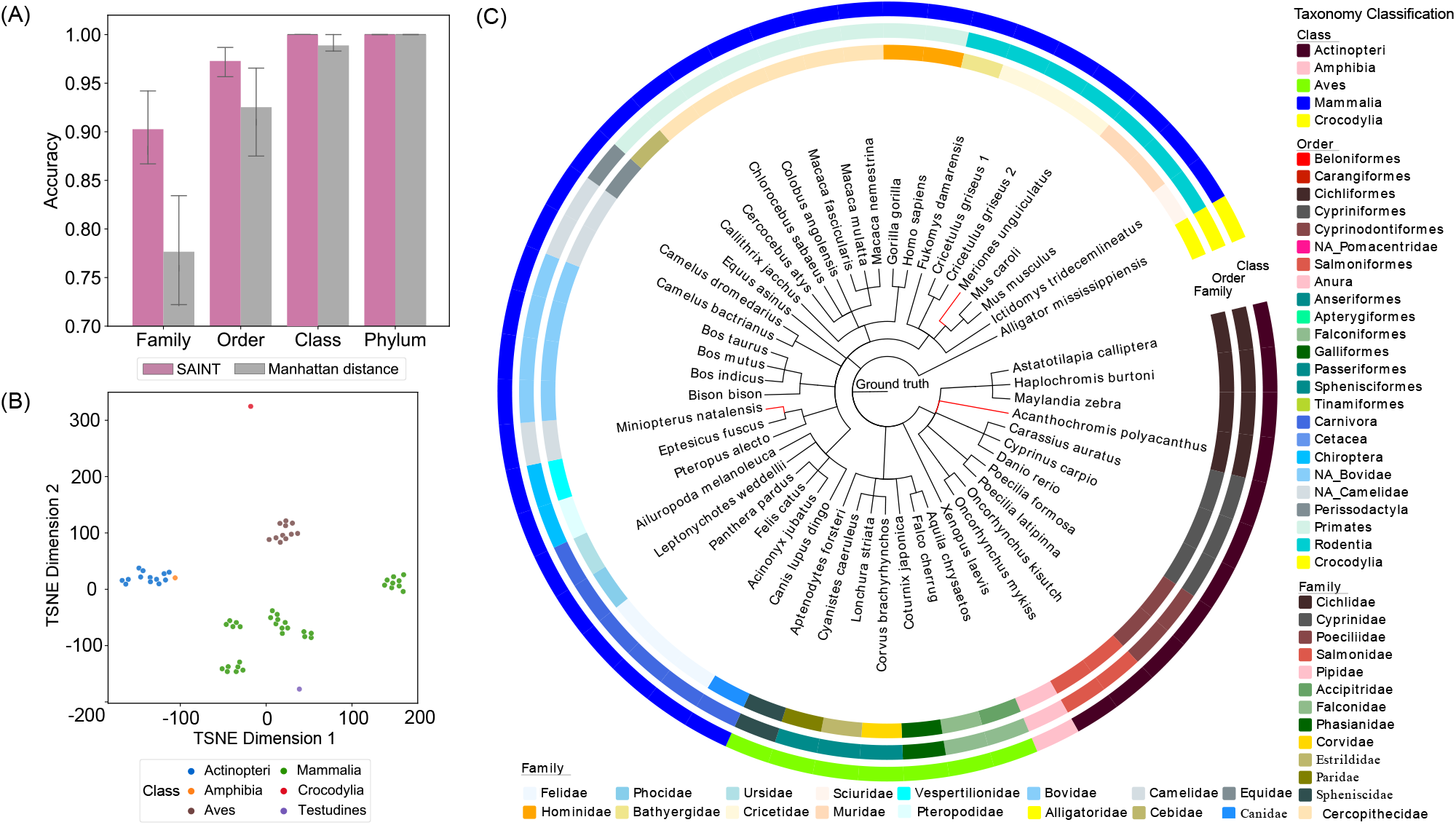
The performance of SAINT in the taxonomy localization task on vertebrate dataset. (A) The prediction accuracy of SAINT and Manhattan distance in terms of different taxonomic level from Genus up to Phylum; (B) The embedding representation learned by SAINT can separate different common vertebrate classes into distinct groups by using t-SNE visualization; (C) The predicted triplets which encode 52 vetebrate reference genomes in the testing set are accurately aligned to the true phytogenetic tree downloaded from NCBI. Note that the branches marked in red indicate the inconsistency between the prediction and the reference phytogeny.

In addition to the prediction accuracy, we also comprehensively evaluated the superior performance of SAINT by treating it as a multi-category classification problem. We used 8 standard measures including precision, recall, F-measure (aka F1 score), homogeneity, completeness, adjusted rand index (ARI), adjusted mutual information (AMI), and normalized mutual information (NMI), to quantitatively assess the performance of SAINT. Their definitions are given in the supplementary material. As shown in Figure 6, SAINT outperformed the Manhattan distance in the majority of cases across various measures. On the bacteria dataset across all taxonomic levels, SAINT achieves an average increase of 6.4%, 4.8%, 5.4%, 2.6%, 3.6%, 9.2%, and 7.2% compared to the Manhattan distance in terms of precision, recall, F1, homogeneity, completeness, ARI, AMI and NMI, respectively. And analogously on the vertebrate dataset, SAINT achieves an average increase of 5.75%, 5%, 6%, 1%, 1.75%, 2%, and 0.5% compared to the Manhattan distance in terms of precision, recall, F1, homogeneity, ARI, AMI, and NMI, respectively

**Fig 6.**
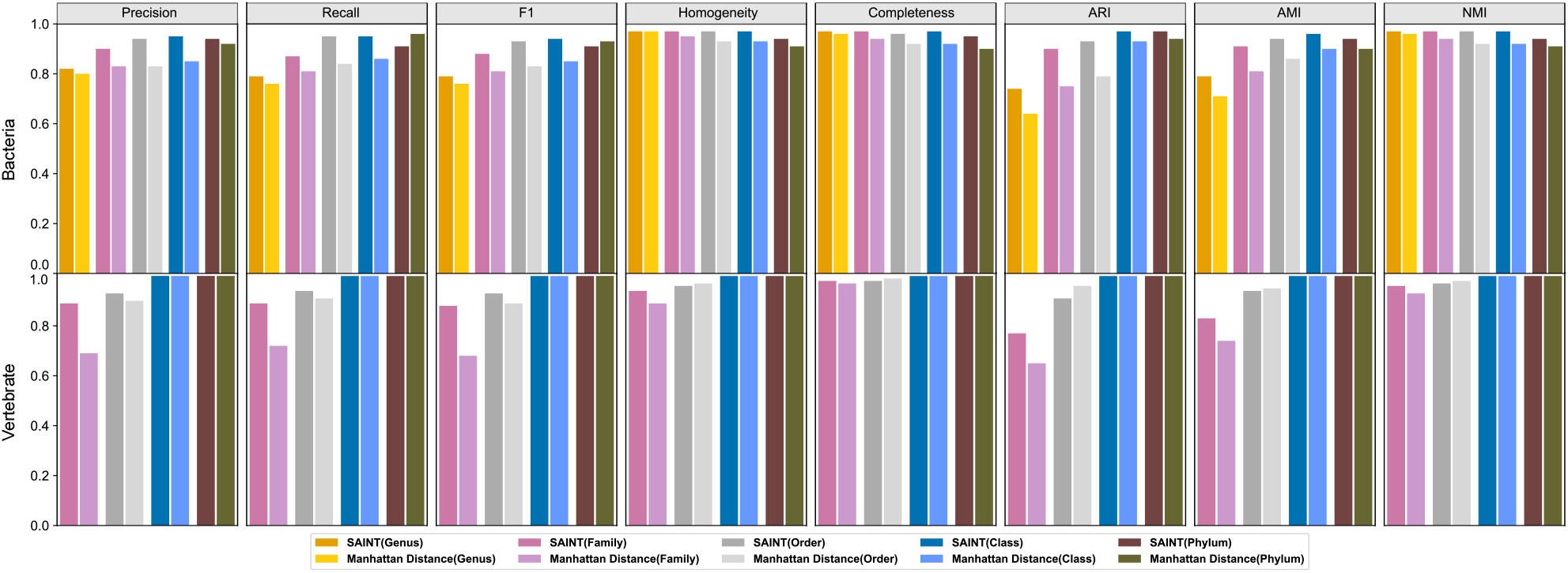
The performance of SAINT treated as a multi-category classification on bacteria and vertebrate datasets, evaluated by multiple measures including precision, recall, F-measure, homogeneity, completeness, adjusted rand index (ARI), adjusted mutual information (AMI), and normalized mutual information (NMI).

The quantitatively good performance of SAINT can also be qualitatively supported, by visualizing the 2-D nonlinear embeddings by using t-SNE (Maaten and Hinton, 2008). For the bacteria dataset shown in Figure 4(B), common phylums, which contain many lower level taxonomic units, such as Proteobacteria, Actinobacteria, and Firmicutes, can be well separated into distinct groups. Meanwhile, relatively rare phylums such as Spirochaetes and Deinococcus-thermus are also distinguishable from others. Analogously, for the vertebrate dataset shown in Figure 5(B), common classes such as Actinopteri, Aves, and Mammalia, can be well separated into distinct groups.

Finally, we aligned the triplets predicted by SAINT with the reference phylogenetic tree from NCBI RefSeq Database (Pruitt *et al*., 2005). To facilitate an unbiased evaluation of SAINT, we consider those triplets whose corresponding sequences are only used in the testing sets (307 out of 1239 bacteria sequences and 52 out of 137 vetebrate sequences in two datasets respectively). For the bacteria dataset shown in Figure 4(C), the 307 bacterial genomes was assigned to 10 phylums. Among the three concentric circles, the outer, middle, and inner circle represent the result of SAINT classification for each genome categorized with respect to Phylum, Order, and Class, respectively. the branches marked in red indicate the wrongly located genomes. Except for negligible bacterial genomes, SAINT accurately locate the majority of genomes into the right place consistently with the reference phylogeney throughout different taxonomic levels. We notice that two misclassifications occur in the Tenericutes phylum, which contains only 5 genomes overall, suggesting that the accuracy of SAINT depends on having enough samples for particular category. The similar conclusion can also be drawn for the vertebrate dataset shown in Figure 5(C).

### 3.4 The effect of various embedding dimensionality

We also performed a parameter sensitivity analysis to investigate how SAINT performs with respect to different embedding dimensionality. As shown in Figure 7, the contrastive loss described in Equation 1 on the testing set exhibit a W-shaped curve. When the embedding dimensionality is small, the contrastive loss drops quickly when the embedding dimensionality increases, until reaching a plateau with training fluctuation. In this paper, we fixed the embedding dimensionality as 100 and potentially, the reported performance can be improved by choosing the dimension corresponding to the first dip.

**Fig 7.**
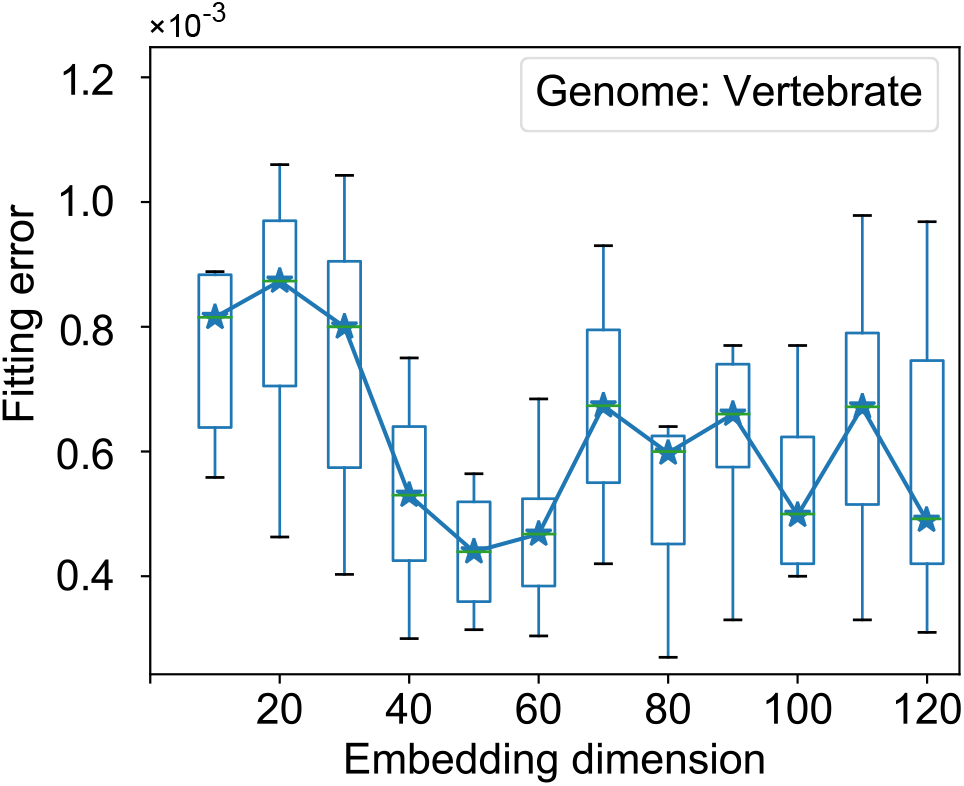
The effect of test error with respect to different embedding dimensionality.

## 4 Conclusion and discussions

In this paper, we developed a novel method, named SAINT, efficient and accurate alignment-free sequence comparison (Figure 1). Compared to existing alignment-free sequence comparison methods, the key novelty of SAINT lies in two aspects: (1) SAINT is a weakly-supervised learning method where the embedding function is learned automatically from the easily-acquired data; (2) SAINT utilizes the non-linear deep learning-based model which potentially better captures the complicated relationship among genome sequences. We have applied SAINT to real-world datasets to demonstrate its empirical utility, both qualitatively and quantitatively. Considering the extensive applicability of alignment-free sequence comparison methods, we expect SAINT to motivate a more extensive set of applications in sequence comparison at large scale.

This work points to several promising directions for future research. First, SAINT requires to represent the input genome sequence as a *k*-mer frequency vector, which potentially lose the interdependent sequence information. In the future, we plan to investigate using other deep neural networks such as convolutional neural networks (CNNs) and recurrent neural networks (RNNs) to process the intact genome sequences as input directly. Second, a desirable property of SAINT model should be transferrable in the sense that the model trained on a particular dataset should be directly applicable to a new dataset, which is vitally important to classify never-before-seen sequences. To this end, we plan to train our model by using a massive collection of sequences from datasets of high volumes, such as the whole NCBI RefSeq Database (Pruitt *et al*., 2005). And finally, aside from achieving a good classification performance, we want SAINT delivers explanation alongside its predictions. For example, if an unseen genome sequence is classified as a coronavirus, then the biologist will want to know which parts of the sequence contribute to this claim. Such explanation capability will enhance the credibility and utility of its predictions for the practitioners.

