## Supplement for "SAINT: automatic taxonomy embedding and categorization by Siamese triplet network"

**Precision,Recall and F1**

Suppose P is the predicted taxonomic clustering results, $\text{TP}_{i},\text{FP}_{i},\text{TN}_{i},\text{FN}_{i}$is calculated for i-th cluster of P，then Precision,Recall and F1 is :

$$P\text{recision}=\sum\frac{\text{TP}_{i}}{\text{TP}_{i}+\text{FP}_{i}}/\left| P \right|$$

$$\text{Recall}=\sum\frac{\text{TP}_{i}}{\text{TP}_{i}+\text{FN}_{i}}/\left| P \right|$$

$$F1=\frac{2\cdot P\text{recision}\cdot R\text{ecall}}{P\text{recision}+R\text{ecall}}$$

**Homogeneity**

Suppose T is the known taxonomic in the Phylogeny tree.The measure homogeneity expects that each cluster contains only the members of a single class. Suppose H(T|P) is the crossentropy of taxonomic types given the cluster P, the homogeneity score is computed by :

$$\text{homogeneity}=\text{1-}\frac{H(\text{T/P})}{H(T)}$$

**Completenes**s

The measure completeness expects that all members of a given class are assigned to the same cluster. The completeness score is computed by :

$$\text{completeness}=\text{1-}\frac{H(\text{T/P})}{H(P)}$$

**ARI**

Suppose n is the total number of samples, $a_{i}$ is the number of samples appearing in the i-th cluster of P, $b_{j}$ is the number of samples appearing in the j-th types of T, and $n_{\text{ij}}$ is the number

of overlaps between the i-th cluster of P and the j-th type and T. ARI is computed as

$$\text{ARI}=\frac{\sum\left( 2^{n_{\text{ij}}} \right)-\left[ \sum\left( 2^{a_{i}} \right)\sum\left( 2^{b_{j}} \right) \right]/\left( 2^{n} \right)}{\frac{1}{2}\left[ \sum\left( 2^{a_{i}} \right)+\sum\left( 2^{b_{j}} \right) \right]-\left[ \sum\left( 2^{a_{i}} \right)\sum\left( 2^{b_{j}} \right) \right]/\left( 2^{n} \right)}$$

**NMI and AMI**

Suppose T is the known taxonomic in the Phylogeny tree, we denote the entropy of P and T as H(P) and H(T), respectively, and the mutual information between them as MI(P,T), $E\left\{ \text{MI}(P,T) \right\}$ is expectation of MI(P,T) . NMI and AMI are computed as:

$$\text{NMI}(P,T)=\frac{\text{MI}(P,T)}{\sqrt{H(P)H(T)}}$$

$$\text{AMI}(P,T)=\frac{\text{MI}(P,T)-E\left\{ \text{MI}(P,T) \right\}}{\text{max}(H(P),H(T))-E\left\{ \text{MI}(P,T) \right\}}$$

For all the measures, larger values (up to 1) mean better performances.
